# Enhanced β-secretase processing of amyloid precursor protein in the skeletal muscle of ALS animal models

**DOI:** 10.1101/352401

**Authors:** Huaqiang Yang

## Abstract

Amyotrophic lateral sclerosis (ALS) is a lethal neurodegenerative disorder primarily characterized by motor neuron degeneration and muscle paralysis. Several studies indicate that pathological changes in the skeletal muscle contribute to disease progression. We report a significant increase of β-secretase processing of amyloid precursor protein (APP) in the skeletal muscle but not the spinal cord or cerebral cortex of hSOD1 (G93A) transgenic ALS mouse models. Enhanced β-secretase processing of APP was manifested by up-regulated expression of βCTF, the 22-kd CTF of APP, and β-secretase processing enzyme, BACE1. Morphological analysis demonstrated that enhanced β-secretase processing of APP mainly occurred in the atrophic myofibers of ALS mice. We also observed a similar change in APP processing in an hSOD1 (G93A) transgenic ALS pig model, suggesting that enhanced β-secretase processing of APP in skeletal muscle may be a common pathological feature of ALS. These findings reveal a selective change in APP processing in skeletal muscle of ALS animal models, and highlight the involvement of aberrant APP processing in ALS pathogenesis.

## INTRODUCTION

Amyotrophic lateral sclerosis (ALS) is a fatal neurodegenerative disease defined by a progressive, selective loss of upper motor neurons (MN) in the motor cortex and lower motor neurons in the spinal cord [1–3]. The degeneration of motor nerves is accompanied by extensive denervation of skeletal muscle and muscular weakness or atrophy, leading to paralysis and ultimately death [1–3]. The mechanisms of MN degeneration in ALS remain unresolved. Abnormal protein aggregation and folding was hypothesized to be one of the major reasons accounting for ALS [4]. Approximately 90% of ALS cases are sporadic, with the remaining 10% of cases being familial. The mutations in the cytoplasmic Cu/Zn superoxide dismutase 1 (SOD1) gene account for approximately 12% of familial ALS cases [5]. A high-copy SOD1 (G93A) transgenic mouse line is commonly used to model ALS and recapitulates much of the pathophysiology of the human disease, including progressive MN degeneration, muscle atrophy, motor deficits, and reduced lifespan [6].

Although MN degeneration in the central nervous system (CNS) is the major pathological characteristic of ALS and has received the most attention, other cell types or tissues beyond the CNS are increasingly accepted to play active roles in ALS [7]. The involvement of skeletal muscle has been the subject of a number of studies conducted on patients and related animal models. Skeletal muscle abnormalities were usually regarded as secondary to the disease mechanisms [8, 9]. However, a large body of evidence demonstrated the importance of skeletal muscle in initiating and contributing to the degeneration of MNs in ALS pathology [7, 10–12]. A study showed skeletal muscle-restricted expression of human SOD1 is sufficient to cause MN degeneration in mice [10]. In contrast, specific neuronal or astrocytic expression of mutant SOD1 in transgenic animals did not result in MN degeneration, clearly negating neurons or astrocytes as the sole mediators of ALS pathology [13–15]. These results demonstrate a causal role of skeletal muscle in ALS and suggest the possibility that skeletal muscle is a key factor for the study of ALS pathogenesis and potential therapy.

Previous studies demonstrated some skeletal muscle-specific molecular changes for ALS [11, 16]. Muscle fiber-specific up-regulation of the amyloid precursor protein (APP) has been shown to occur in both SOD1 mutant mice, as well as in human ALS cases before the appearance of clinical abnormalities [11]. Moreover, genetic ablation of APP significantly improved multiple disease parameters and ameliorated disease progression in ALS mouse models [17]. These findings further indicate that the pathological events that occur locally in the skeletal muscle contribute to the CNS impairment, and reversal of these changes can alleviate the corresponding lesion in the CNS. APP and its processing products are usually involved in Alzheimer’s disease (AD) [18]. Mutations in APP are associated with the familial form of AD. β-amyloid protein (Aβ), a cleavage product of APP, accumulates at high levels in the brain in AD, which constitutes the typical pathological feature of this disease. Overproduction or aggregation of Aβ in the brain is a primary cause of AD [19]. In this process, APP is cleaved by β-secretase to generate amyloidogenic carboxyterminal fragments (βCTP), which is further cleaved by γ-secretase to generate Aβ [18, 19]. The similar changes in APP expression or processing in the affected tissues in AD or ALS suggest interesting parallels among these neurodegenerative disorders. Although elevated levels of APP in ALS muscle have been reported, whether and how APP processing in ALS skeletal muscle changes are still unclear. Because APP cleavage products are usually involved in the pathological process leading to neuronal degeneration, we here examined the changes in the expression of APP β-secretase processing products and enzyme in the skeletal muscle of mouse and pig models of ALS.

## RESULTS

### β-secretase processing of APP was enhanced in skeletal muscle but not the CNS in ALS mouse models

The hindlimb skeletal muscle (gastrocnemius), lumbar spinal cord, and cerebral cortex from end-stage (120 days) SOD1 (G93A) transgenic mice and 120-day-old wild-type (WT) littermates were collected and subjected to Western blot to examine the expression levels of APP and APP cleavage products. Staining with the Y188 antibody specific to the carboxy-terminal end of APP showed a 10-kd and a 15-kd fragment in both WT and ALS mice. The amount of the fragment approximately 15 kd in size was significantly higher in the skeletal muscle but not the brain or spinal cord in the ALS mice than in the WT mice. In addition, we observed significantly increased expression of the 22-kd fragment in the skeletal muscle of ALS mice (Figure 1). The mouse Aβ-specific antibody detected 15- and 22-kd fragments but failed to detect a 10-kd fragment (Figure 2). These results suggested that the 10-kd and 15-kd fragments were αCTF and βCTP fragments, respectively, and the 22-kd fragment might be a longer Aβ-containing APP CTF.

**Figure 1.**
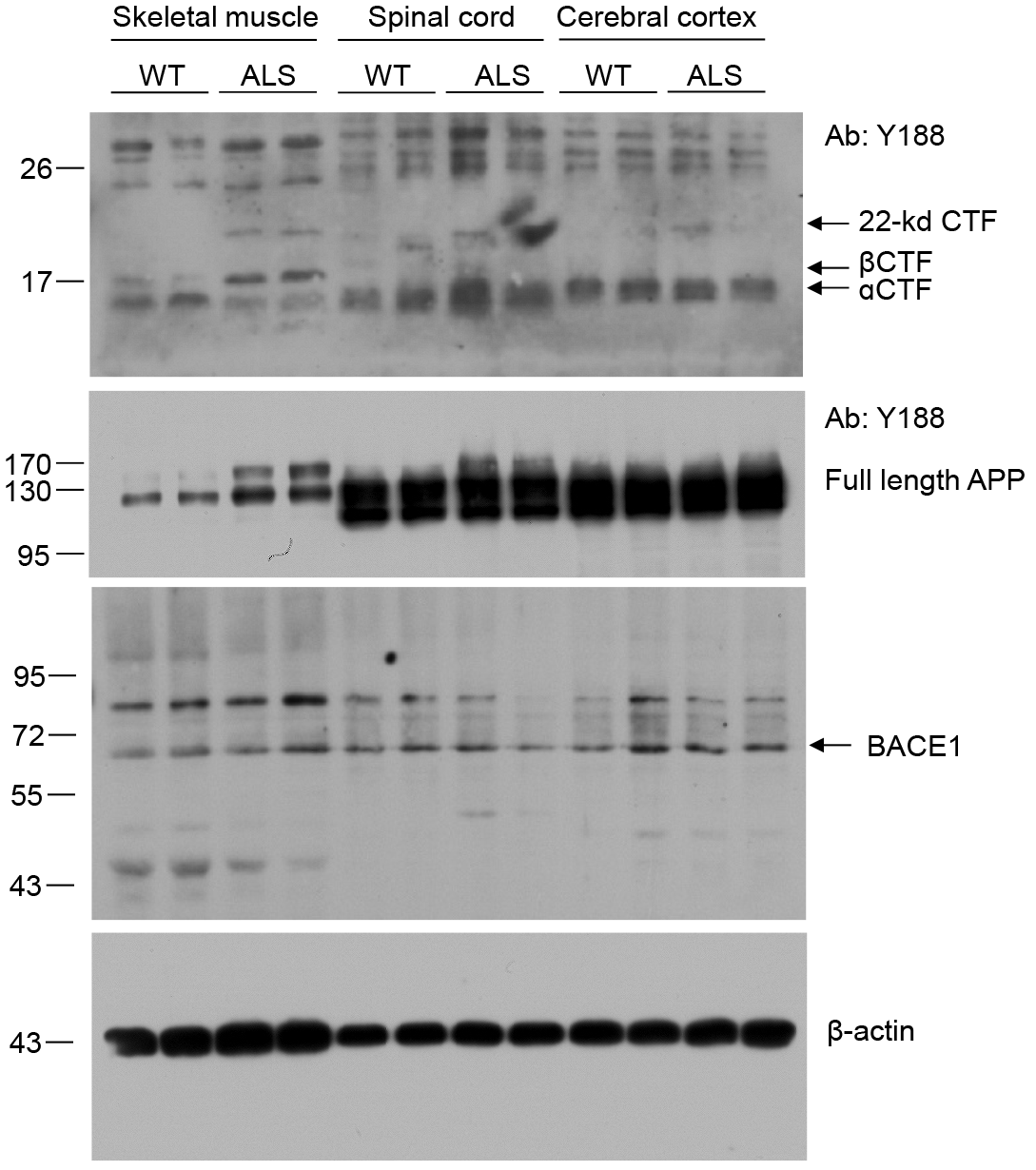
Selectively elevated β-secretase processing of APP in skeletal muscle of ALS mice. Elevated levels of β-CTF and 22-kd CTF were observed in the skeletal muscle but not in the spinal cord and cerebral cortex of ALS mice compared with those of WT mice. Meanwhile, selectively elevated expression of full-length APP and BACE1 was detected in skeletal muscle of ALS mice. Two mice were used as representatives for each group to examine the expression levels of the indicated proteins.

**Figure 2.**
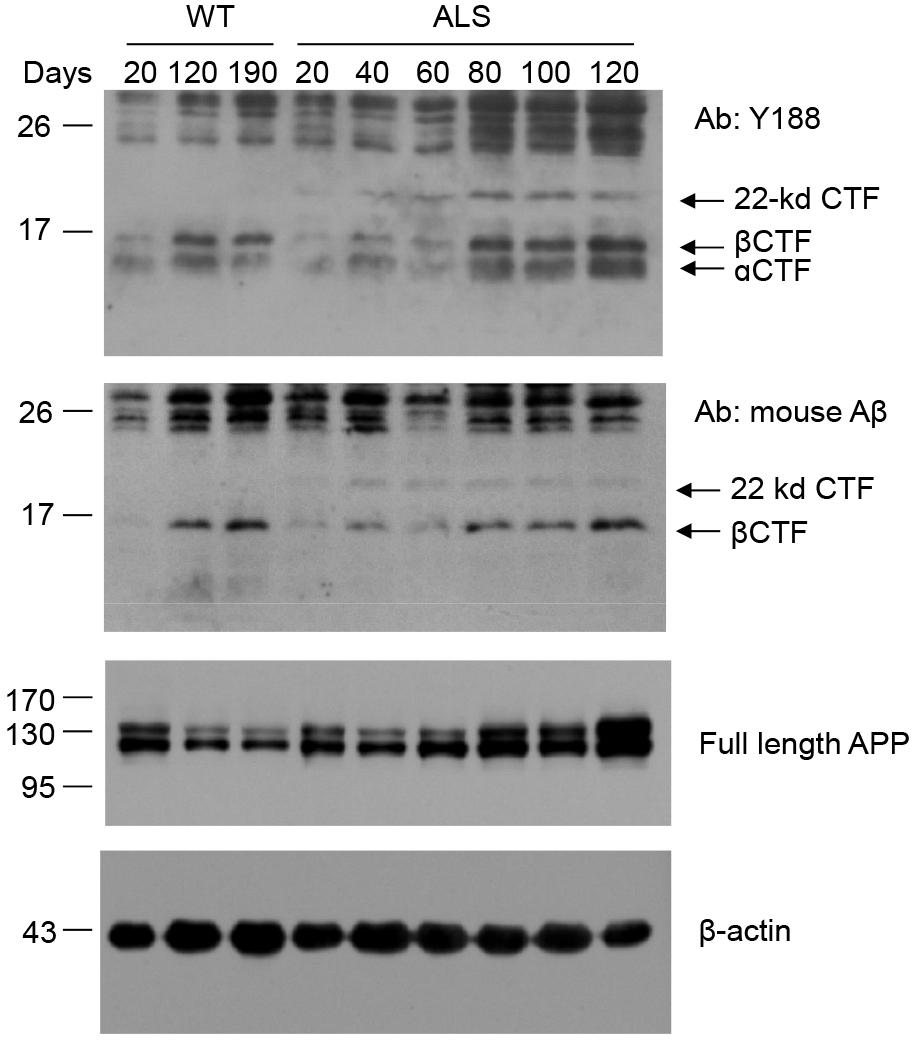
Time-course study of β-secretase processing of APP in the skeletal muscle of ALS mice. The skeletal muscles of ALS mice at 20, 40, 60, 80, 100, and 120 days of age were subjected to Western blot to detect the change in β-secretase processing with time. The β-CTF and 22-kd CTF were observed in the skeletal muscle of ALS mice as early as 20 days of age, and the expression levels gradually increased over the 120-day period. β-CTF expression also increased over time in WT controls, but its expression level was significantly lower than that of the age-matched ALS mice. No 22-kd CTF was observed in WT control animals at any time point. An antibody specific to mouse Aβ also detected an increase in β-CTF and 22-kd CTF over time. Expression of full-length APP in the skeletal muscle of ALS mice gradually increased starting at 40 days of age.

We next detected the β-secretase processing products of APP in the skeletal muscle of ALS mice at different days of age. We found that the 22-kd fragment could be observed in the skeletal muscle of ALS mice as early as 20 days of age and persisted until death, whereas no 22-kd fragment was detected in WT mice at any time point. The βCTP fragment was significantly up-regulated in the skeletal muscle of ALS mice starting at 80 days of age and then increased with age. Although the amount of βCTP fragment also increased with aging in the skeletal muscle of WT mice, there was significantly less protein in the skeletal muscle of WT mice than in the age-matched ALS mice at the same time points (Figure 2).

The skeletal muscles from different parts of the body were collected from the ALS mice at end-stage disease. In addition to gastrocnemius, the tibialis anterior, quadriceps femoris, gluteus maximus, triceps brachii, trapezius, and pectoralis major all showed up-regulated expression of βCTP and 22-kd fragments compared with those in WT mice (Figure 3). The results indicated that β-secretase processing of APP was enhanced in skeletal muscles all over the body.

**Figure 3.**
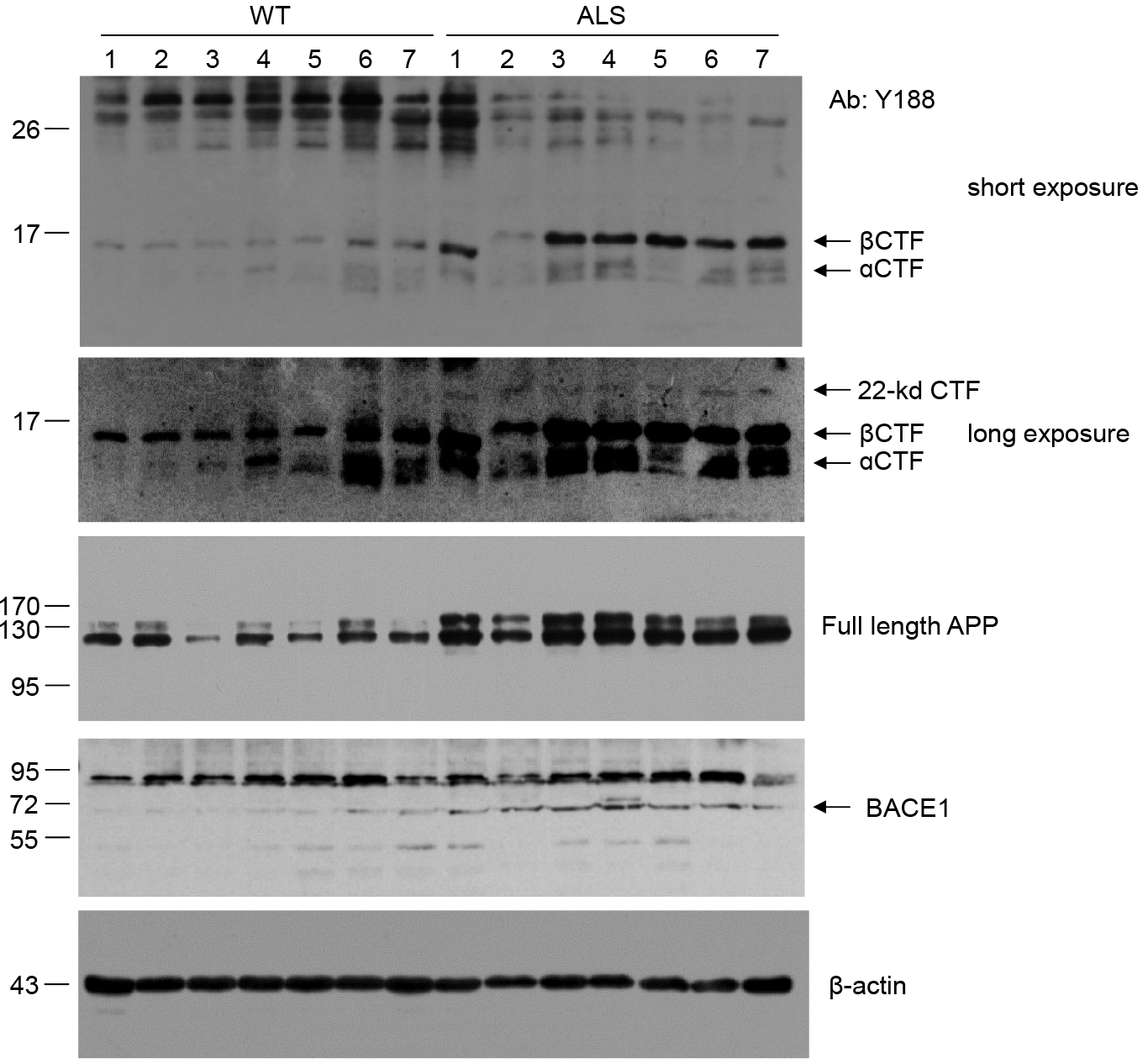
Elevated β-secretase processing of APP in various skeletal muscles throughout the body of ALS mice. β-CTF and 22-kd CTF expression showed significant up-regulation in skeletal muscles from different parts of the body of ALS mice. The top two images were the same blot under short and long exposures, respectively, to reveal the α-CTF, β-CTF, and 22-kd CTF expression. Y188 and BACE1 blotting showed up-regulation of full-length APP and BACE1, respectively, in multiple types of skeletal muscle from ALS mice. 1, gastrocnemius; 2, tibialis anterior; 3, quadriceps femoris. 4, gluteus maximus; 5, triceps brachii; 6, trapezius; 7, pectoralis major.

### Enhanced expression of BACE1 in skeletal muscle of ALS mice

Since β-site APP cleaving enzyme 1 (BACE1) is the major β-secretase [20], we detected the expression level of BACE1 in the skeletal muscle, lumbar spinal cord, and cerebral cortex of ALS and WT mice. Western blot assay demonstrated that BACE1 expression levels were specifically up-regulated in skeletal muscle but not the brain or spinal cord in the ALS mice (Figure 1). We also observed up-regulated BACE1 expression in various skeletal muscles throughout the body of ALS mice (Figure 2). The enhanced BACE1 level corresponded to the enhanced β-secretase processing of APP, suggesting that BACE1 plays a part in elevating the β-secretase processing of APP in ALS skeletal muscle.

### Enhanced β-secretase processing of APP occurred in atrophic muscle fibers

We sectioned the gastrocnemius of the end-stage WT and ALS mice and stained the cross-section of the muscle fibers using mouse Aβ and BACE1 antibodies. Immunohistochemistry (IHC) demonstrated significantly enhanced Aβ staining, which represents the Aβ or Aβ-containing APP CTF, in ALS muscle fibers compared with that in WT muscle fibers (Figure 4A). Most positive fibers had a smaller cross-section area than negative ones in the ALS muscle, indicating that enhanced β-secretase processing of APP mainly occurs in the atrophic muscle fibers (Figure 4B).

**Figure 4.**
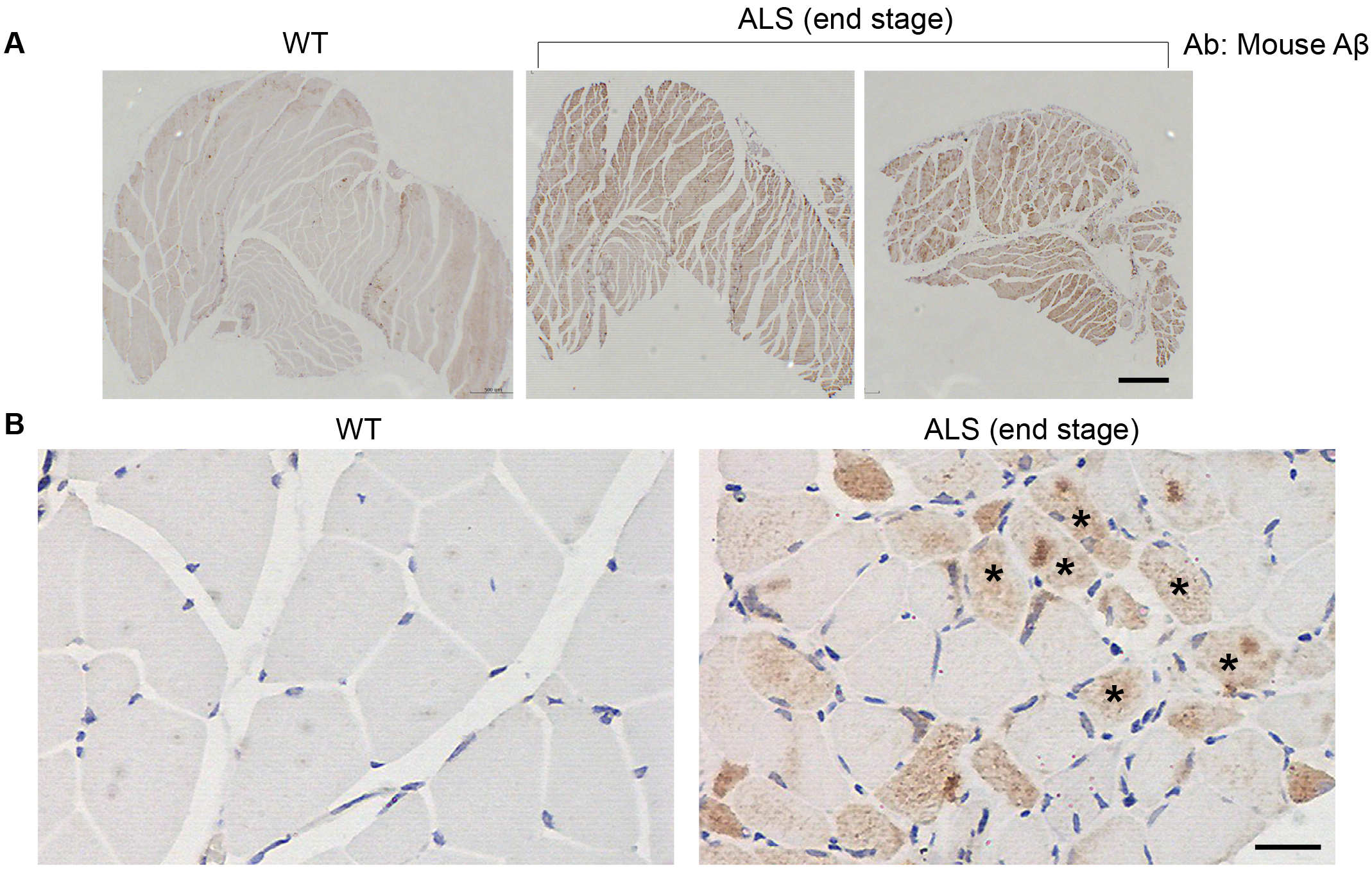
IHC showed significantly elevated APP β-secretase processing in skeletal muscle of ALS mice. **(A)** Staining of the gastrocnemius with a mouse Aβ antibody showed a significant up-regulation in the APP β-cleaved product in ALS mice. **(B)** The positive muscle fibers exhibited greater morphological atrophy (asterisk) than the negative ones. Scale bars: 500 μm (upper panel), 20 μm (lower panel).

A similar staining pattern was observed for BACE1. BACE1 IHC showed significantly enhanced BACE1 staining in ALS muscle compared with that in WT muscle, and the positive signal was mainly distributed in the atrophic muscle fibers (Figure 5). IHC results further confirmed the above Western blot results, showing that β-secretase processing of APP was greatly enhanced in the skeletal muscle of ALS mice.

**Figure 5.**
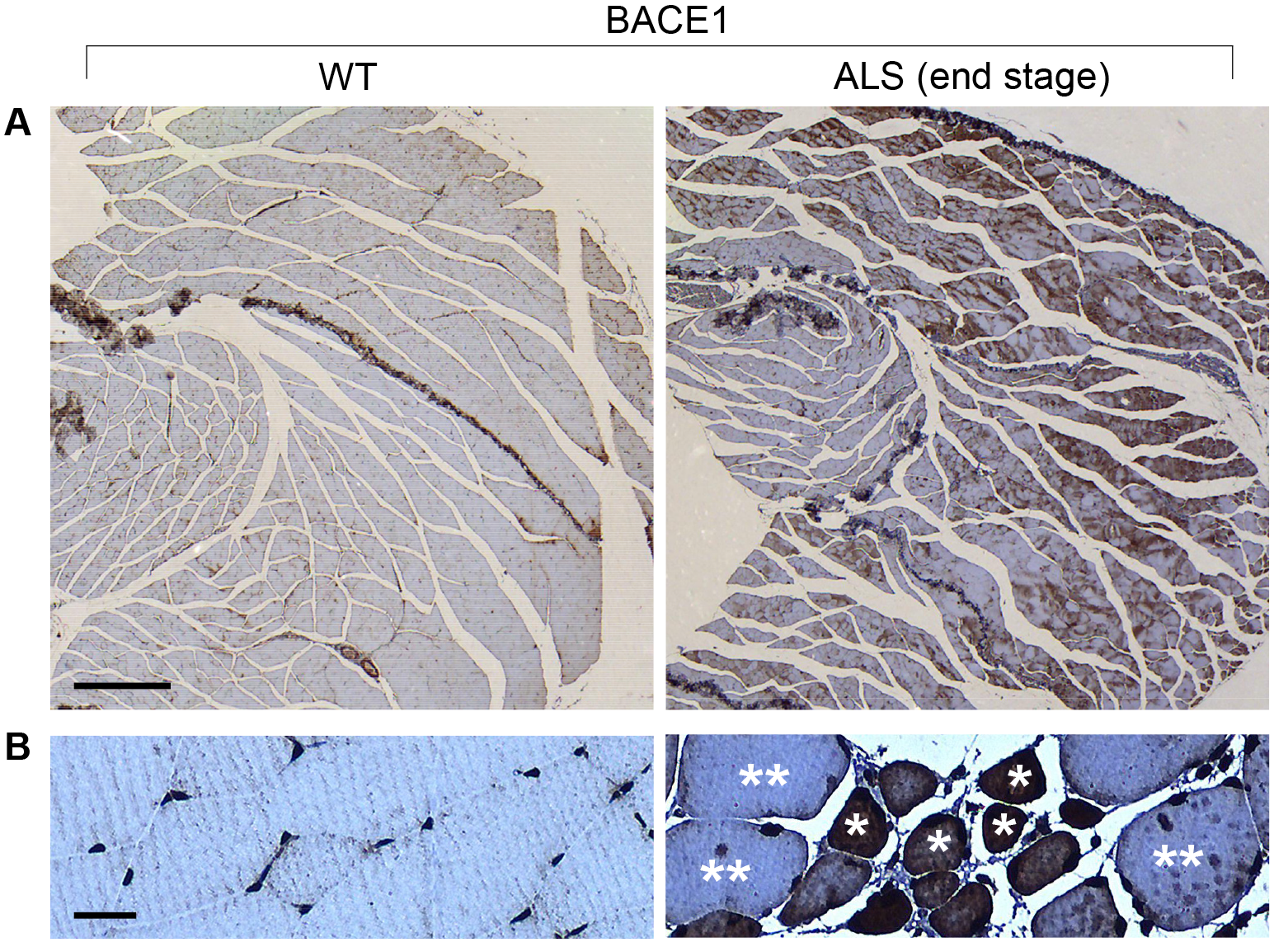
IHC showed significantly elevated BACE1 expression in skeletal muscle of ALS mice. **(A)**Significant up-regulation of BACE1 in the gastrocnemius of ALS mice. **(B)** BACE1-positive staining was mainly localized to atrophied muscle fibers (asterisk). Double asterisks indicate negatively stained fibers without morphological atrophy. Scale bars: 400 μm (upper panel), 20 μm (lower panel).

### SOD1 mutant ALS pig models showed elevated β-secretase processing of APP in skeletal muscle

We have established an ALS pig model harboring a human SOD1 transgene with a G93A mutation [21]. To investigate the changes in APP processing in the pig model, the gastrocnemii of ALS and WT pigs were assayed for the related protein levels at the age of 8 months when the motor deficit was significant in the ALS pigs. Western blots showed significantly greater βCTF and BACE1 expression in the muscle of ALS pigs than in that of WT pigs, suggesting that the β-secretase processing of APP was enhanced in ALS pig models (Figure 6A). However, unlike in the mouse model, the level of full-length APP was not elevated in the muscle of ALS pigs, nor could the 22-kd fragment be observed in ALS pig muscle. IHC using antibody 2454 recognizing human/pig Aβ demonstrated significantly greater immunoreactivity in the skeletal muscle fibers of ALS pigs than in those of WT pigs (Figure 6B). Since IHC staining is reportedly negative for full-length APP in skeletal muscle fibers of WT animals [17], and no up-regulated expression of full-length APP was observed in ALS pig muscle by Western blot, the positive signal may represent the up-regulated expression of the Aβ-containing CTF. We further confirmed these results by staining the skeletal muscle fibers using another antibody, MOAB-2, which specifically recognizes human/pig Aβ but not full-length APP. A similar staining pattern was observed in the skeletal muscle fibers of ALS pigs, showing significantly greater β-secretase processing of APP than in the WT animals (Figure 6C). BACE1 IHC also demonstrated greater expression levels in ALS pig muscles than in WT muscles. Moreover, βCTF and BACE1 positive staining was mainly limited to atrophic muscle fibers (Figure 6D). Double immunofluorescence labeling showed co-localization of βCTF and BACE1 immunoreactivity in the ALS muscle fibers (Figure 7).

**Figure 6.**
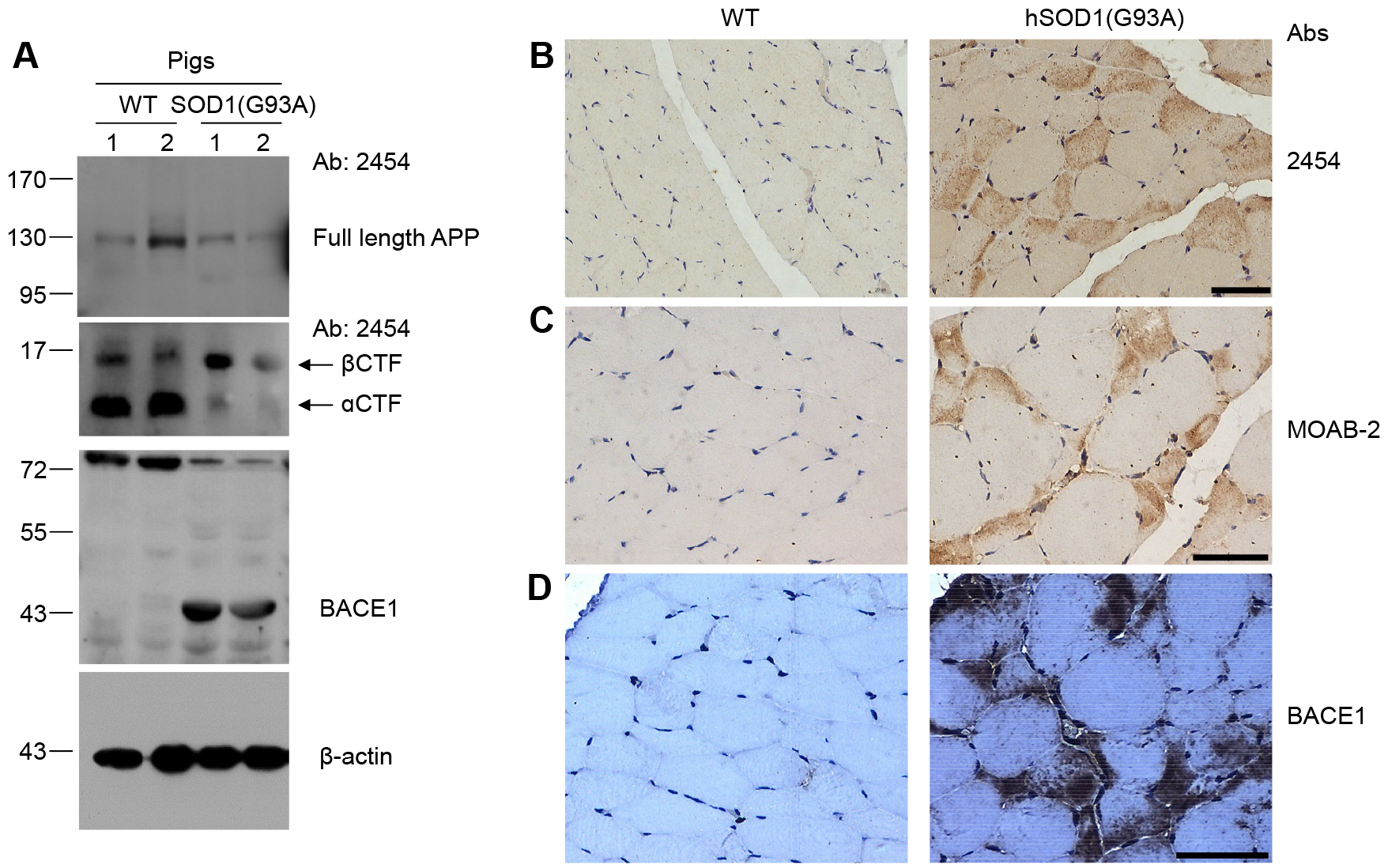
Elevated β-secretase processing of APP in skeletal muscle of ALS pig models. **(A)**Western blot assays showed elevated β-CTF and BACE1 expression in ALS pigs. The expression level of full-length APP showed no significant change between the ALS and WT pigs. **(B)** IHC using antibody 2454 showed positive staining in skeletal muscle fibers of ALS pigs. **(C)** IHC with the antibody MOAB-2, which does not recognize full-length APP, indicated that the APP β-cleaved product but not full-length APP was up-regulated in the muscle of ALS pigs. **(D)** The skeletal muscle of ALS pigs showed up-regulated BACE1 expression compared with that of WT pigs. Scale bars: 50 μm.

**Figure 7.**
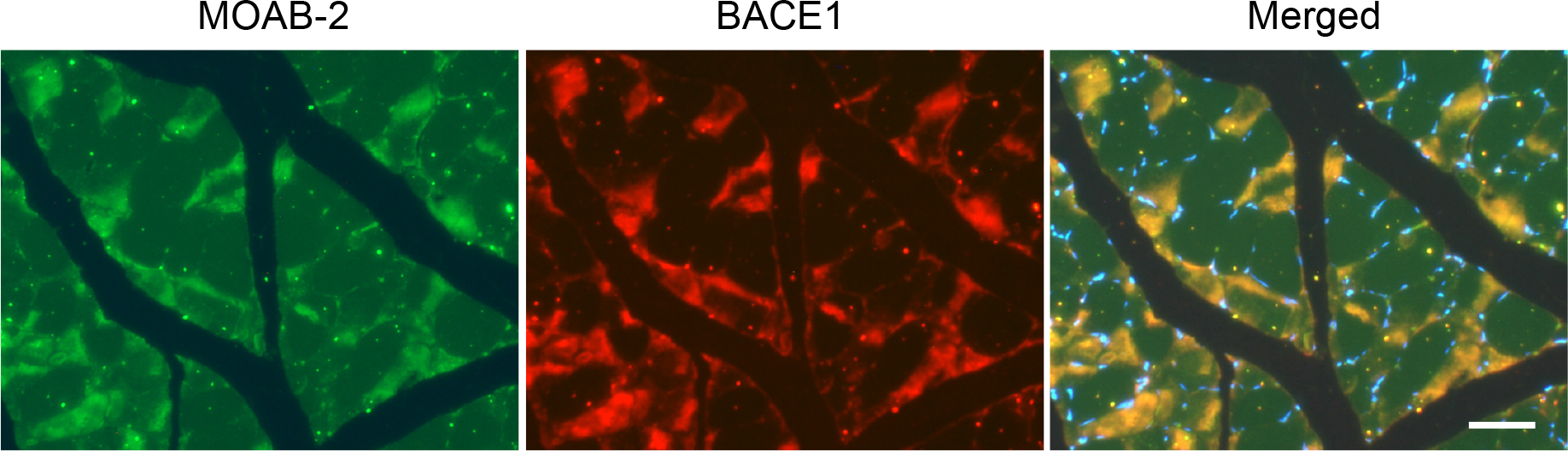
Co-localization of β-CTF and BACE1 in skeletal muscle of ALS pigs. Double immunofluorescence labelling of β-CTF (green) and BACE1 (red) demonstrated their co-localization in atrophied skeletal muscle fibers. Scale bar: 50 μm.

## DISCUSSION

This study provides the first demonstration that β-secretase processing of APP is significantly elevated in the skeletal muscle of ALS animal models. These observations extend the previous report of increased APP expression in the skeletal muscle of ALS patients and mouse models. The elevation in APP processing preceded the onset of disease signs in ALS mice by as early as 20 days, implying that APP processing is important for triggering other physical signs, including neuronal lesion and skeletal muscle paralysis. In addition to βCTF, we reproducibly detected a 22-kd CTF of APP in the skeletal muscle of ALS mice but not WT mice. The antibody recognizing Aβ also detected this fragment, indicating that the 22-kd CTF is a longer amyloidogenic processing product of APP. Previous studies reported the existence of the Aβ-containing 22-kd CTF in human platelets [22], brain microvessels [23], transfected 293 cells [24, 25], and human umbilical vein endothelial cells [24]. The enhancement of βCTF and 22-kd CTF levels in skeletal muscle is an evident early pathological event in ALS mouse models, hence could be potential therapeutic targets.

The roles of APP processing or cleavage products in the pathogenesis of ALS have been implicated in several studies [26–29]. SOD1 mutant ALS mouse models treated intraventricularly with a monoclonal antibody that blocks the β-secretase cleavage site on APP at a presymptomatic stage (70 or 90 days) resulted in a decrease in the levels of soluble APP-β and a delay in disease onset and deterioration [27]. In another study, over-expression of βCTF led to the earlier onset of motor symptoms associated with ALS [28]. These reports combined with our observations demonstrate that APP cleavage products might contribute to the degeneration in ALS, and early inhibition of the APP process may ameliorate disease progression. Because the disease in skeletal muscle is an initiating causal pathological process in ALS, not merely secondary neurogenic atrophy, and elevation of APP processing is a muscle-specific early event in ALS pathology, related studies should focus on skeletal muscle rather than only on the CNS. A strategy for an APP processing intervention in skeletal muscle needs to be developed to evaluate its therapeutic potential for ALS.

In this study, the change in β-secretase processing of APP is manifested by an enhanced expression of βCTF and BACE1. However, we have been unsuccessful in detecting Aβ in skeletal muscle by immunoblot or ELISA (data not shown). Whether the failure to detect Aβ peptide reflects a high turnover or inefficient production of Aβ peptide in skeletal muscle is presently unknown. Similarly, soluble Aβ_x−40_ and Aβ_x−42_ in muscle tissue from WT and ALS mice were reportedly undetectable by ELISA [17]. Indeed, the detection of Aβ peptide (Aβ monomer) in skeletal muscle with immunoblot seems to be difficult at present. Sporadic inclusion-body myositis (s-IBM) is a muscle disease characterized by accumulation of Aβ within muscle fibers. However, immunoblot assay did not detect Aβ monomers in s-IBM muscle [29]. In another study on transgenic mice over-expressing βCTF, a high level of βCTF was detected, but Aβ was difficult to detect with an immunoblot assay in the skeletal muscle and many other peripheral tissues of the transgenic mice [30]. These findings indicate the need for the development of a more sensitive approach to detect the Aβ monomer in skeletal muscle. Furthermore, the Aβ level in neurons of the lumber spinal cord of ALS patients and mouse models was reportedly enhanced [17, 31], whereas our result clearly showed no enhancement of APP β-secretase processing in the spinal cord and brain compared with that in WT animals, raising concerns that increased Aβ deposition may not necessarily be related to the increased β-secretase processing of APP.

In regard of the accumulated βCTF as we observed, the toxicity of βCTF accumulation has been confirmed in other studies [32, 33]. A triple transgenic AD mouse model shows an early accumulation of βCTF in the brain, and the anatomical lesions in the brain is related with βCTF levels [32]. Another paper documents that βCTF, rather than Aβ, causes memory deficits in a mouse model of dementia [33]. In the present work, we observed a similar early and age-dependent accumulation of βCTF in the skeletal muscle of ALS models, further suggesting its implication as an early and key contributor of the neurodegenerative process taking place in the animal models.

We also observed a similar change in APP processing in skeletal muscle of pig models harboring mutant SOD1 expression. The pig models have been well characterized in pathological and behavioral features as an ideal large animal model for ALS [21]. Due to the anatomical, physiological, and genetic similarities between humans and pigs, pigs are considered to be better animal models than rodents to resemble human disease; therefore, the outcome from pig models might be easily translated to the clinic [34]. The elevated APP β-secretase processing in skeletal muscles of both mouse and pig models implies that the aberrant APP processing in skeletal muscle is a general pathological feature of ALS. Taken together, amyloidogenic APP CTF and BACE1 were selectively elevated in the skeletal muscles of ALS mouse and pig models. APP β-secretase processing in the skeletal muscle may be a potential target for the development of a disease-modifying approach for ALS.

## MATERIALS AND METHODS

### Animals

Transgenic mice harboring a high copy number of the hSOD1 (G93A) transgene [B6SJL-Tg(SOD1-G93A)1Gur/J], as reported by Gurney et al., were used in this study [6]. The breeding pairs were purchased from The Jackson Laboratory, and the hemizygous transgenic progeny were maintained by mating the transgenic males with F1 hybrid wild-type females. The progeny were genotyped by PCR using genomic DNA isolated from the tail after birth as previously described [6]. The transgenic pigs used in this study harbored an hSOD1 (G93A) transgene and were reported previously [21]. All experiments were performed in accordance with the guidelines for animal experiments of the Department of Science and Technology of Guangdong Province, China, and approved by the animal research committee of the Guangzhou Institutes of Biomedicine and Health.

### Western blot

For Western blotting, tissue samples were homogenized in RIPA buffer and centrifuged at 16,000 g at 4 °C for 30 minutes. Supernatants were collected and the protein concentration was quantified using a Pierce BCA protein assay kit (Thermo Scientific). Lysates were subjected to SDS polyacrylamide gel electrophoresis, followed by immunoblotting onto a polyvinylidene fluoride membrane (Millipore). Membranes were blocked with 5% non-fat dry milk in TBST for 2 hours and incubated overnight at 4 °C with the primary antibodies shown in Table 1. After the membranes were washed with TBST, they were further incubated with a secondary HRP-conjugated goat anti-mouse or rabbit IgG antibody for 60 minutes at room temperature and imaged using a SuperSignal West Pico enhanced chemiluminescence kit (Thermo Scientific). The actin antibody (sc-47778, Santa Cruz) was used as a loading control.

**Table 1.** Antibodies used in this study

| Antibody | Recognized targets | Reactivity | Applications | Source |
| --- | --- | --- | --- | --- |
| Y188 | Full-length APP,<br>βCTF, αCTF | Mouse | WB | Abcam |
| Mouse Aβ | βCTF | Mouse | WB, IHC | Abcam |
| 2454 | βCTF | Pig | WB, IHC | Cell signaling |
| MOAB-2 | βCTF, does not<br>recognize full-length<br>APP | Pig | IHC | Cayman |
| BACE1 | BACE1 | Mouse,<br>pig | WB, IHC | Proteintech |

### Immunohistochemistry

The skeletal muscles were fixed with 4% buffered paraformaldehyde for at least 48 h. The tissues were then dissected, embedded in paraffin wax, cut in cross-section, and mounted on a slide. Sections were deparaffinized in xylene and rehydrated in a graded series of ethanol. Antigen retrieval was performed in a 1 M citrate buffer (pH = 6.0) bath at 90-100 °C for 20 min. Sections were blocked with 5% goat serum in PBST for 1 hour at room temperature, stained with the indicated primary antibodies overnight at 4 °C, and visualized using a VECTASTAIN ABC detection systems (Vector Laboratories). Nuclei were counterstained with hematoxylin. Finally, sections were dehydrated in graded ethanol, cleared in xylene, and then coverslipped with Permount.

### Double immunofluorescence

Double immunofluorescence staining was used to examine the co-distribution of APP β-secretase cleavage products and BACE1 in the same sample. After antigen retrieval, tissue sections were incubated with MOAB-2 and BACE1 primary antibodies, followed by incubation with Alexa Fluor 488-labeled goat anti-mouse IgG and Alexa Fluor 594-labeled goat anti-rabbit IgG secondary antibodies (Invitrogen). DAPI was used to label nuclear DNA. The slides were analyzed by fluorescence microscopy (Olympus).

## CONFLICTS OF INTEREST

The authors declare no conflicts of interest.

## FUNDING

This work was partially supported by grants from the National Natural Science Foundation of China (31772555).

